# A Sensitized ENU Mutagenesis Screen for Thrombosis Modifiers Identifies Suppressor Variants in Non-mutagenized Parental Generations Due to Antithrombotic Selective Pressures

**DOI:** 10.64898/2026.01.25.701592

**Authors:** Marisa A. Brake, Audrey C. Cleuren, Sydney Torres, Martijn A. van der Ent, Guojing Zhu, Justin Kulchycki, Tyler M. Parsons, Caitlin D. Schneider, Adrianna M. Jurek, Kailey MacFadyen, Amy E. Siebert, David R. Siemieniak, Andrew E. Timms, David R. Beier, Randal J. Westrick

## Abstract

Thrombosis is a leading cause of morbidity and mortality. We used a mouse forward genetic ENU screen to identify genomic variants that suppress *F5*^L/L^ *Tfpi*^+/-^ lethal thrombosis. Surviving *F5*^L/L^ *Tfpi*^+/-^ mice from our *M*odifier of *F*actor *5 L*eiden 16 (*MF5L16*) ENU line were subjected to whole-genome sequencing analysis. This revealed that instead of an ENU-induced mutation, four mutations introduced from our *F5*^L/L^ breeding stock were responsible for survival, which we named *sMF5L1-4* for *s*pontaneous *M*odifier of *F*actor *5 L*eiden. In our colony, *F5*^L/L^ female breeders carrying all four *sMF5L* mutations produced more litters and offspring than breeders with three or fewer mutations (p<0.006). Genotyping of 13 additional *MF5L* lines demonstrated that the four *sMF5L* mutations were present in all lines and were consistently associated with survival. Of these four mutations, a single G to A intergenic variant on Chromosome 18 (Chr18^A^, *sMF5L4*), was most significantly associated with survival (p=0.003), with ∼15% penetrance for conferring the survival phenotype. Furthermore, platelet aggregation was significantly reduced in Chr18^A^ mice, suggesting an additional mechanism by which Chr18^A^ could suppress lethal thrombosis. Comparative transcriptomics analysis of livers from Chr18^A^ mice versus wildtype littermate controls revealed a small number of differentially expressed genes both known and unknown to affect thrombosis. In summary, we have identified four variants exerting a significant selective breeding advantage along with antithrombotic effects. Superimposing our mutagenesis screen on a selective background illustrates the interplay of natural strain background variants and *de novo* ENU mutations in suppressing *F5*^L/L^ *Tfpi*^+/-^ lethal thrombosis.

## Introduction

Venous thromboembolism (VTE) is a leading cause of morbidity and mortality, causing significant health issues and affecting quality of life for afflicted patients^1^. VTE is complex as it is associated with and influenced by other medical disorders such as obesity, recent surgery, prolonged immobility, cancer, chronic kidney diseases, and infectious diseases^2^. Additionally, VTE exhibits a high degree of heritability, with genetics accounting for approximately 50% of the VTE risk. Well-documented genetic susceptibility variants such as Factor V Leiden (FVL) and gene-level deficiencies in antithrombin, Protein C, and Protein S can predispose carriers to developing VTE^3^.

Large scale human genome-wide association studies (GWAS) including whole exome and whole genome sequencing have identified numerous loci associated with VTE. Interestingly, many of the genes linked to the phenotype are outside of the canonical coagulation pathway^4–8^. While the use of proteomics has enabled the direct investigation of protein levels and their association to VTE risk^8^, most of the highly associated GWAS variants are within noncoding regions with unknown regulatory functions (e.g., intronic, upstream, downstream, or intergenic variants)^7,11^. More recent evidence suggests that complex traits can be modified by several small effect variants that interact and work together to contribute to disease, with many of those variants residing in noncoding regions^9,10^. *In-silico* GWAS meta-analysis and transcriptome-wide association studies have revealed that many significant noncoding variants are found in enhancer regions and histone-marked sites in blood and liver tissue^12^. These regions could provide a more global layer of regulatory control of hemostasis in addition to the more localized control of gene expression due to individual promoter elements. This could suggest that noncoding variants are more prevalent and consequential in VTE than previously thought. Thus, a remaining challenge is to establish direct genotype-phenotype relationships for loci impacting VTE^13^.

FVL is an unusual mutation because it is common and exerts a large effect on VTE risk^14^. However, FVL also exhibits incomplete penetrance and variable expressivity for the development of thrombosis^15^, suggesting a genetic susceptibility trait that can be influenced by genetic modifiers^3^. Mice with the Factor V Leiden mutation (*F5*^L^) display a mild prothrombotic phenotype that closely mirrors the human disease^16^, and *F5*^L^ homozygosity (*F5*^L/L^) combined with hemizygosity for tissue factor pathway inhibitor (*Tfpi*^+/-^) results in synthetic perinatal lethality^17^. To identify genetic modifiers of FVL, we used the *F5*^L/L^ *Tfpi*^+/-^ thrombotic phenotype to perform a sensitized mouse ENU (N-ethyl-N-nitrosourea) mutagenesis screen for novel VTE modifiers that restore normal survival. For our initial work, we identified 16 surviving mouse lines, termed *M*odifier of *F*actor *5 L*eiden 1-16 (*MF5L1-16*)^18^. Using whole exome sequencing, we identified one major VTE modifier mutation in the *Actr2* gene for the *MF5L12* line. However, there were no clear modifiers in the remaining 15 lines^18^.

For the present studies, we used whole genome sequencing of mice from our *MF5L16* ENU line to identify additional mutations within noncoding regions that suppresses the lethal thrombotic phenotype of *F5*^L/L^ *Tfpi*^+/-^ mice. We determined that these mutations arose spontaneously in the *F5*^L^ breeding stock used to generate mice for the mutagenesis screen, and we named these mutations *sMF5L1-4* for *s*pontaneous *M*odifier of *F*actor *5 L*eiden. The *sMF5L* mutations conferred significant survival and propagation advantages to the *F5*^L/L^ mice. Of these four mutations, a single G to A intergenic variant on Chromosome 18 (Chr18^A^, *sMF5L4*) had the most significant effect, in addition to reducing platelet reactivity and driving transcriptional changes in genes with links to coagulation. Our results demonstrate that the partial lethality observed in *F5*^L/L^ homozygous mice is sufficient to drive evolution in a breeding colony. Our sensitized ENU mutagenesis screen was superimposed upon this evolving *F5*^L/L^ background, with the *sMF5L4* modifiers contributing to the suppression of thrombosis across many of our *MF5L* lines.

## Results

### MF5L16 Mice Exhibit a Coagulation Defect

Whole exome sequencing analysis for the highly penetrant line of *F5*^L/L^ *Tfpi*^+/-^ mice (*MF5L16*, 73.7% penetrance of survival) failed to identify a single ENU induced mutation that segregated with a majority of surviving mice^18^. To examine this line for possible alterations in blood coagulation, we performed the Prothrombin Time (PT) assay, which detects defects in fibrinogen, thrombin, FV, FVII, and FX^19^, and is sensitive to *Tfpi* differences^20^. As expected, *Tfpi*^+/-^mice trended towards a shorter clotting time (p=0.065) and *F5*^+/L^ *Tfpi*^+/-^ had a significantly shorter time compared to C57BL/6J (B6) (p=0.039; Fig. 1). In contrast, the *MF5L16 F5*^L/L^ *Tfpi*^+/-^ mice exhibited a significantly longer time compared to *Tfpi*^+/-^ and *F5*^+/L^ *Tfpi*^+/-^ controls (p<0.0001; Fig. 1), resulting in a PT similar to B6. These PT assay results suggest that *MF5L16 F5*^L/L^ *Tfpi*^+/-^ mice possess a genetic variant (or variants) that could modify the extrinsic plasma protein activities as measured by the PT, thereby providing biochemical evidence for a mechanism by which the *MF5L16* thrombosis suppressor(s) rescues the lethal phenotype.

**Figure 1:**
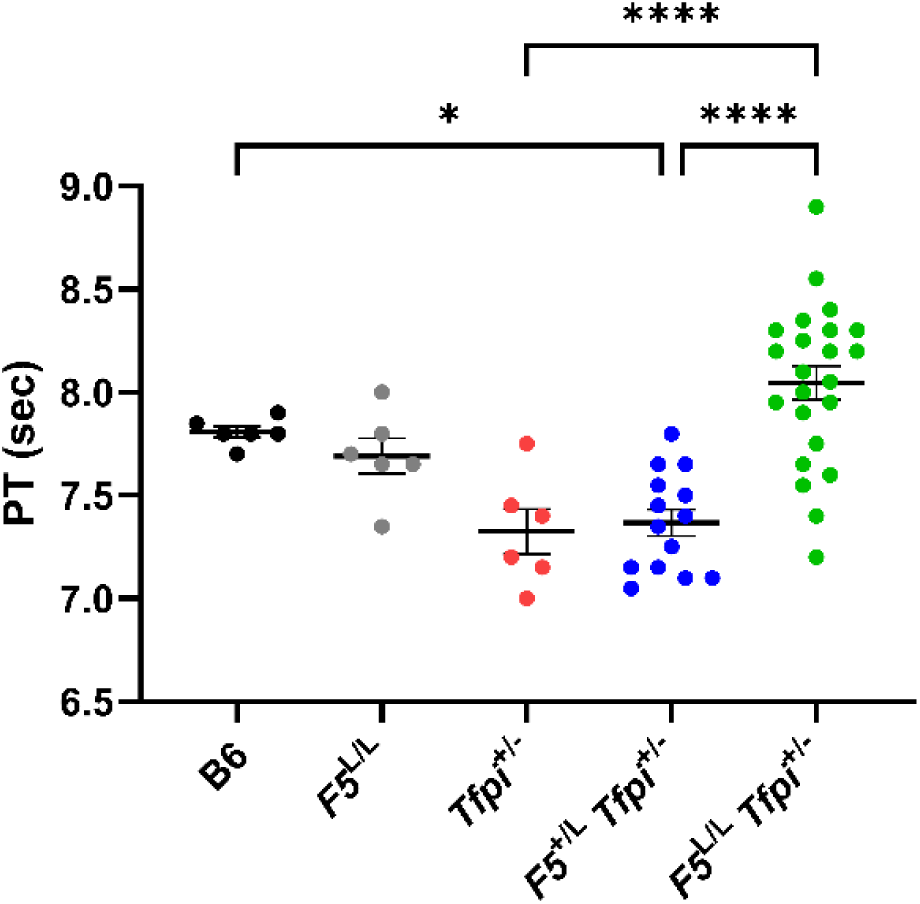
Prothrombin Time (PT) of *MF5L16* Mice. *F5*^L/L^ *Tfpi*^+/-^ mice display a longer time to clot compared to *Tfpi*^+/-^ and *F5*^+/L^ *Tfpi*^+/-^ mice. *Tfpi*^+/-^ and *F5*^+/L^ *Tfpi*^+/-^ mice have a shorter time to clot compared to B6 and *F5*^L/L^ mice. Each data point represents a biological replicate. Mean ± SEM. One-way ANOVA, *p<0.05, ****p<0.0001

### Whole Genome Sequencing Analysis Identifies sMF5L Mutations Introduced from Non-mutagenized Parental Breeding Stock Associated with Increased Survival

Whole genome sequencing (WGS) was performed on four *MF5L16 F5*^L/L^ *Tfpi*^+/-^ mice (276-794 days of age), resulting in the identification of 329 variants. Only two of these variants were exonic, within the genes *Dhrsx* and *Peg10* (Supplemental Table S1). The remaining top candidates were selected based on quality by depth ≥2^21^, resulting in 32 candidate ENU mutations (Supplemental Table S1). Each of these candidates was validated by Sanger sequencing of the four mice subjected to whole genome sequencing as well as parental B6 and 129S1/SvImJ (129S1) controls. Twenty-two of the candidates, including the two exonic variants, were classified as false-positive calls as they were not present in the original WGS mice. Interestingly, we identified three large, previously uncharacterized genomic insertions/deletions between B6 and the reference genome as well as between the B6 and 129S1 mouse strains, by Sanger sequencing analysis of the candidate point mutations (Supplemental Fig. S1). The remaining 10 candidates were identified as valid mutations, with three introduced from the *Tfpi*^-^ background and seven introduced into *MF5L16* by the non-mutagenized *F5*^L/L^ breeders.

Spontaneous mutations develop gradually over multiple generations of breeding, both in wildtype mice and in genetically engineered mice at an increased rate^22–24^. Previous studies have demonstrated that spontaneous mutations can have a significant effect on phenotypes and thrombosis suppression^25–28^. Therefore, the seven mutations introduced from the *F5*^L/L^ breeders were analyzed in all 136 *MF5L16* mice. Four mutations were significantly associated with survival while three were not (Fig. 2, Supplemental Fig. S2), with a G to A intergenic variant on Chromosome 18 (Chr18^A^, *sMF5L4*) at position 63,103,082, exhibiting the most significant effect on survival (p=0.003, Fig. 2A). The other three significant mutations were an A to G mutation in a genomic region with a predicted gene (Gm42243) and two predicted lncRNAs (ENSMUSG00000097726 and ENSMUSG00000127837) on Chromosome 5 (Chr5^G^, *sMF5L1*) at position 29,059,210 (p=0.04, Fig. 2B), a C to T mutation in the intron of *4933400C23Rik* on Chromosome 9 (Chr9^T^, *sMF5L2*) at position 92,629,622 (p=0.02, Fig. 2C), and an A to G mutation in the intron of *Zfp131* on Chromosome 13 (Chr13^G^, *sMF5L3*) at position 120,238,235 (p=0.05, Fig. 2D). Analysis of interactions between these four spontaneously arising mutations demonstrated that inheritance of all four mutations is most significantly associated with survival of *F5*^L/L^ *Tfpi*^+/-^ *MF5L16* mice (p=2×10^-5^).

**Figure 2:**
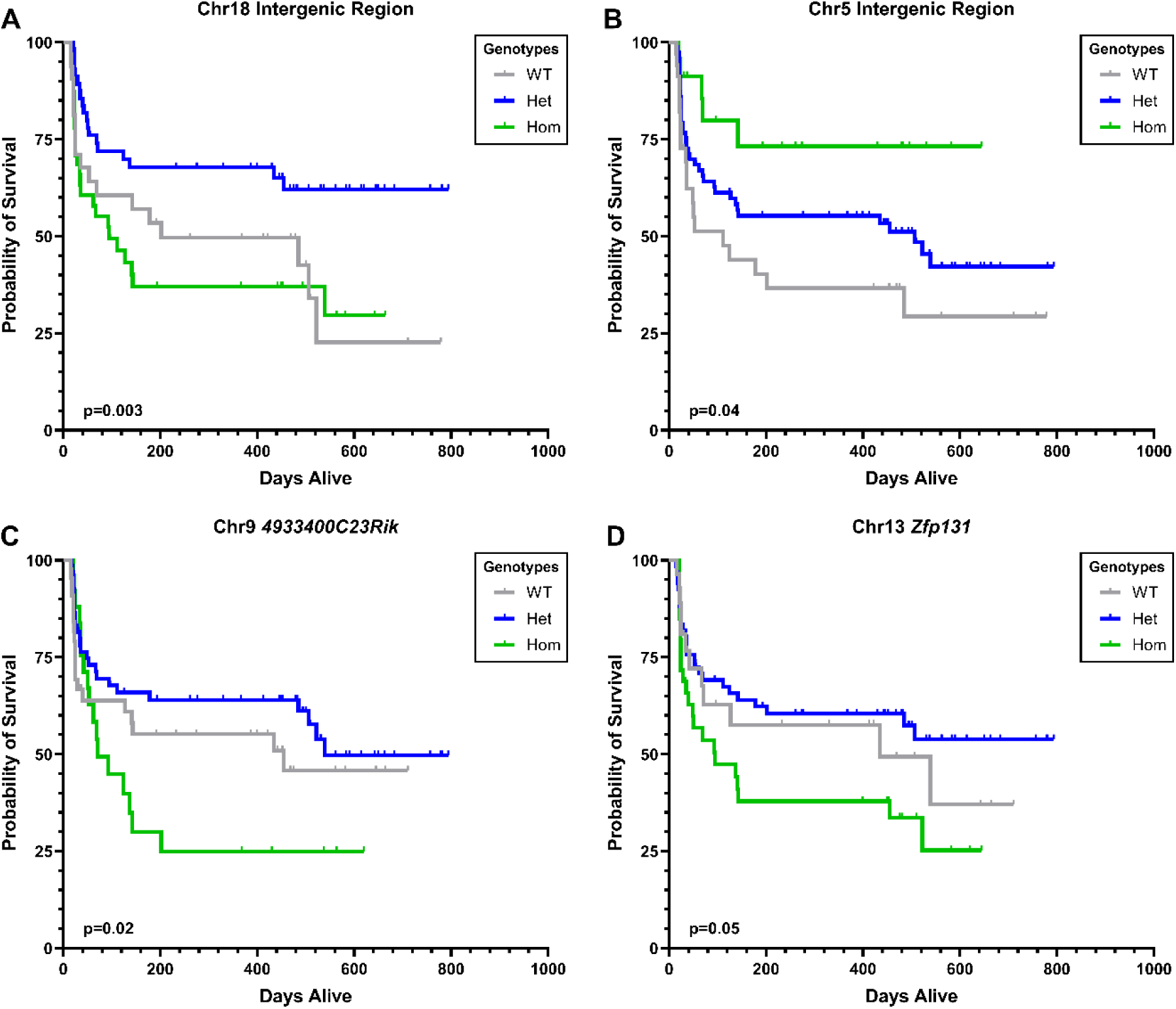
Four Introduced Mutations are Significantly Associated with Survival. Significant Kaplan-Meier survival curves of *F5*^L/L^ *Tfpi*^+/-^ mice from *MF5L16*. **(A)** Curve between WT (n=32), Het (n=58), and Hom (n=47) for the Chr18 intergenic region mutation. **(B)** Curve between WT (n=34), Het (n=79), and Hom (n=24) for the Chr5 intergenic region mutation. **(C)** Curve between WT (n=44), Het (n=67), and Hom (n=26) for the Chr9 intronic mutation in *4933400C23Rik*. **(D)** Curve between WT (n=28), Het (n=67), and Hom (n=42) for the Chr13 intronic mutation in *Zfp131*. WT=wildtype; Het=heterozygous for mutation; Hom=homozygous for mutation

### The sMF5L Mutations Increase Fecundity of F5^L/L^ Mice

To better understand the evolutionary basis underlying the persistence of these mutations, we investigated the effects of the four *sMF5L* mutations on fitness and reproduction by analyzing historical DNA samples of offspring from 240 *F5*^L/L^ breeders genotyped for each *sMF5L* from our *F5*^L/L^ colony. Breeding pairs were grouped into four categories based on their genotypes for each *sMF5L* mutation: 1) both breeders wildtype; 2) one breeder wildtype and one heterozygous; 3) one breeder wildtype and one homozygous; and 4) both breeders heterozygous or homozygous for the mutation. For each *sMF5L* mutation, the average number of litters and offspring produced by the *F5*^L/L^ breeders with the *sMF5L* mutations was greater than those in which both breeders were wildtype (Supplemental Table S2). When all four *sMF5L* mutations were carried by the *F5*^L/L^ breeders, significantly more litters (4.93 ± 0.23, p=0.0004) and offspring (20.9 ± 0.88, p=0.006) were produced compared to *F5*^L/L^ breeders with three or fewer mutations present (3.47 ± 0.32 and 16.03 ± 1.43, respectively) (Figure 3A; Supplemental Table S2). The greater number of offspring from the *F5*^L/L^ breeders with all four *sMF5L* mutations suggests these mutations lead to improved fecundity and provide an evolutionary mechanism for the persistence of these mutations in our *F5*^L/L^ colony.

**Figure 3:**
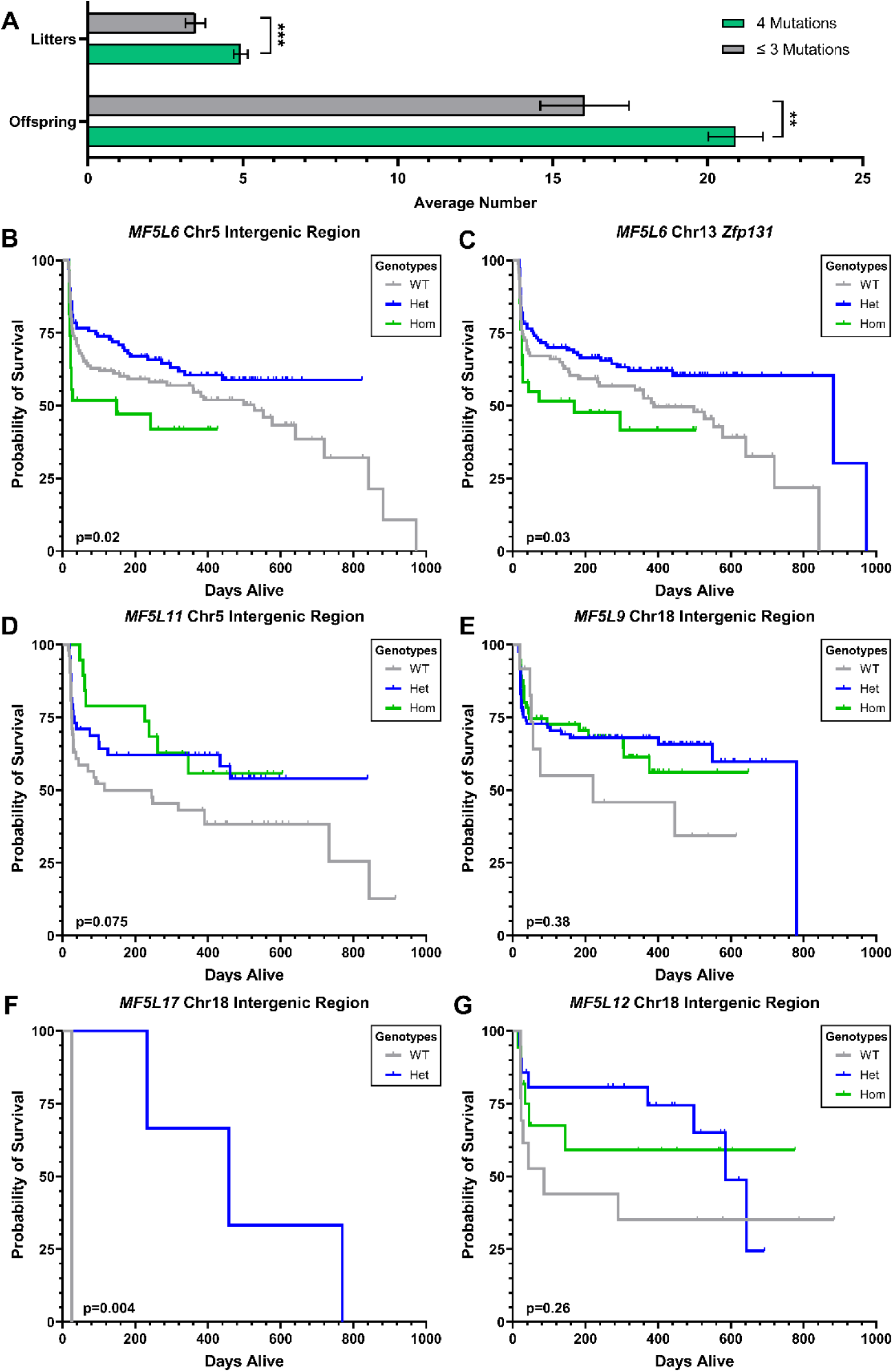
Introduced Mutations are Significantly Associated with Fecundity and Survival in Other *MF5L* Lines. **(A)** Average number of litters and offspring in the *F5*^L/L^ breeders carrying all four (n=58 matings) and three or less (n=36 matings) of the introduced mutations. Mean ± SEM. Student’s t-test, **p<0.01, ***p<0.001. **(B)** Survival curve between WT (n=126), Het (n=112), and Hom (n=27) for the Chr5 intergenic region mutation and **(C)** between WT (n=96), Het (n=134), and Hom (n=34) for the Chr13 intronic mutation in *Zfp131* in *MF5L6* mice. **(D)** Survival curve between WT (n=53), Het (n=49), and Hom (n=19) for the Chr5 intergenic region mutation in *MF5L11* mice. **(E)** Survival curve between WT (n=12), Het (n=89), and Hom (n=56) for the Chr18 intergenic region mutation in *MF5L9* mice. **(F)** Survival curve between WT (n=2) and Het (n=2) for the Chr18 intergenic region mutation in *MF5L17* mice. **(G)** Survival curve between WT (n=13), Het (n=21), and Hom (n=18) for the Chr18 intergenic region mutation in *MF5L12* mice. Interactions between Chr18 and *Actr2* in *MF5L12* demonstrate both mutations are associated with survival. WT=wildtype; Het=heterozygous for mutation; Hom=homozygous for mutation

### The sMF5L Mutations are Present in All MF5L Lines and are Associated with Survival

All additional 1,197 mice from 13 other *MF5L* lines were genotyped for the four *sMF5L* mutations, which were present to varying degrees in every *MF5L* line. Knowing the contribution of these mutations within each line will enable other mutations conferring survival to be identified. However, all of these lines have a B6/129S1 mixed background. Since the four *sMF5L* mutations arose in the B6 *F5*^L/L^ mice, *MF5L* mice that were crossed to 129S1 *F5*^L/L^ parental mice should not have these mutations. *F5*^L/L^ *Tfpi*^+/-^ mice on the 129S1 background may still harbor a thrombosuppressive mutation, either inheriting an ENU-induced or (more rarely) a spontaneous mutation arising in the 129S1 *F5*^L/L^ mice. Therefore, the survival analyses for lines with large numbers of 129S1 mixed *F5*^L/L^ *Tfpi*^+/-^ mice were performed with and without the mice with a large contribution of 129S1 ancestry.

In the *MF5L6* line, we previously identified a linkage peak on Chromosome 3, which encompasses the gene *F3*, but no coding mutations were identified in that region^18^. Analysis of 265 B6 *F5*^L/L^ *Tfpi*^+/-^ mice from this line revealed that both Chr5^G^ and Chr13^G^ are significantly associated with survival (p=0.02, Fig. 3B and p=0.03, Fig. 3C, respectively). In 121 *F5*^L/L^ *Tfpi*^+/-^mice from the *MF5L11* line, Chr5^G^ heterozygosity and homozygosity trended toward increased survival (p=0.075; Fig. 3D). In 157 B6 *F5*^L/L^ *Tfpi*^+/-^ mice from the *MF5L9* line, Chr18^A^ heterozygosity and homozygosity also appeared to increase survival, although this did not reach significance (p=0.38; Fig. 3E). In a small *MF5L* line not previously reported (*MF5L17*), the founding *F5*^L/L^ *Tfpi*^+/-^ mouse only produced four *F5*^L/L^ *Tfpi*^+/-^ out of 15 total offspring. The Chr18^A^ mutation was present in all *F5*^L/L^ *Tfpi*^+/-^ mice except for the one that died at 25 days. Thus, Chr18^A^ is significantly associated with survival (p=0.004; Fig. 3F). The penetrance of Chr18^A^ in suppressing lethality in *MF5L17* is 53.3%, which is the strongest manifestation of this allele in any *MF5L* line.

We previously identified a mutation in *Actr2* as a potent thrombosuppressor of *F5*^L/L^ *Tfpi*^+/-^lethal thrombosis in *MF5L12*, however, *Actr2* did not account for 100% of the survival^18^. Analysis of the *sMF5L* mutations in 51 *MF5L12* mice demonstrated that Chr18^A^ alone trends toward survival (p=0.26; Fig. 3G). Furthermore, Chr18^A^ has a strong interaction with the *Actr2* mutation, suggesting that co-segregation of both mutations is most significantly associated with survival (p=0.001).

### In silico Determination of Regulatory Function of the sMF5L Mutations

The four *sMF5L* mutations reside in genomic regions without known regulatory elements or highly conserved regions. Therefore, we determined the predicted regulatory function of these mutations in four different tissue types compared to the wildtype allele by a deep learning convolutional network, DeepSEA (Table 1)^28,29^. DeepSEA is a tool to accurately predict epigenetic events such as transcription factor binding, DNase I sensitivities, and histone marks in a variety of cell types to determine the probability that a given sequence is a regulatory feature. The megakaryoblastic cancer cell line, CMK, was used to represent megakaryocytes, and CD34+ mobilized cells were used to represent hematopoietic stem cells.

**Table 1:** Predicted Regulatory Functional of the Introduced Mutations.

|  | Allele | Hepatocyte | Aortic Smooth Muscle Cell | Megakaryoblastic Cancer Cell Line (CMK) | CD34+ Mobilized Cells |
| --- | --- | --- | --- | --- | --- |
| Chr18 | + | 0.00 % | 0.00 % | 14.9 % | 0.00 % |
|  | A | 1.14 % | 5.52 % | -7.71 % | -9.56 % |
| Chr5 | + | 0.00 % | 0.00 % | 0.00 % | 0.00 % |
|  | G | -1.08 % | 4.18 % | 0.33 % | 2.57 % |
| Chr9 | + | 0.00 % | 0.00 % | 0.00 % | 0.00 % |
|  | T | 3.26 % | -0.12 % | -6.33 % | -6.17 % |
| Chr13 | + | -7.47 % | -17.6 % | -3.76 % | -4.22 % |
|  | G | 8.75 % | 4.55 % | 18.9 % | 15.3 % |

The Chr18^A^ mutation had an increased chance of regulatory function in hepatocytes and aortic smooth muscle cells, but a decreased chance of functionality in the CMK and CD34+ cells, suggesting this genomic region could contain a cell type-specific expression motif. The Chr5^G^ mutation had a decreased chance of functionality in hepatocytes and an increased functionality in the other three cell types. In contrast, the Chr9^T^ mutation had an increased chance of functionality in hepatocytes and a decreased functionality in the other three cell types. Lastly, the Chr13^G^ mutation had an increased chance of functionality in all tissue types. Although the percent functionality was low, the mutations were predicted to instigate a change in the predicted regulatory function of those genomic sequences, suggesting these mutations could be putative global or cell-specific regulatory elements/regions.

### Chr18^A^ Suppresses F5^L/L^ Tfpi^+/-^ Lethality

We tested the effect of the single Chr18^A^ variant on *F5*^L/L^ *Tfpi*^+/-^ mouse survival by breeding *F5*^+/L^ *Tfpi*^+/-^ Chr18^+/A^ triple heterozygous mice to *F5*^L/L^ Chr18^A/A^ mice. While a genotype ratio of ∼1:8 for *F5*^L/L^ *Tfpi*^+/-^ Chr18^+/A^ mice was expected, only two *F5*^L/L^ *Tfpi*^+/-^ Chr18^+/A^ mice were produced out of a total of 109 offspring: one female and one male. The female mouse survived to 466 days, was phenotypically normal, and reproduced normally. She was crossed with a *F5*^L/L^ Chr18^+/+^ mouse and produced 20 pups in total. Only one of her 20 offspring was *F5*^L/L^ *Tfpi*^+/-^, but it did not carry Chr18^A^ and died before weaning. The male rescue mouse survived 173 days and also reproduced normally. He was crossed with a *F5*^L/L^ Chr18^+/+^ mouse and produced 10 pups, none with the *F5*^L/L^ *Tfpi*^+/-^ genotype. This male had to be euthanized because of seizures, and autopsy of the brain showed clots and deterioration of the left frontal lobe (Supplemental Fig. S3). If the Chr18^A^ variant was fully penetrant for the survival phenotype, we would expect ∼13 *F5*^L/L^*Tfpi*^+/-^ Chr18^+/A^ mice to have been produced out of 109 total offspring. Taken together, these results suggest that Chr18^A^ accounts for ∼15% of the survival penetrance and that the three other *sMF5L* mutations, Chr5^G^, Chr9^T^, and Chr13^G^, may work together with Chr18^A^ to improve survival.

### Chr18^A^ Mice Exhibit Normal Coagulation but Have a Platelet Functional Defect

To further determine if the Chr18^A^ mutation affects coagulation, we performed the PT and activated Partial Thromboplastin Time (aPTT) assays, a tail bleeding time, and measured plasma factor V levels. Chr18^A^ conferred similar PT and aPTT time to clot as wildtypes, and there were no differences in the total bleeding time between mice with or without the Chr18^A^ mutation (Supplemental Fig. S4A-C). Circulating FV levels were quantified by ELISA, and we observed no significant differences in FV between the three genotypes (Supplemental Fig. S4D).

To determine if the Chr18^A^ mice had a platelet defect, we performed ADP and collagen-induced whole blood platelet aggregation. The area under the curve (U) and aggregation (AU) demonstrated no difference in Chr18^A/A^ mice compared to wildtype controls with either agonist, however, the Chr18^+/A^ mice had significantly reduced area under the curve (U) with collagen (p=0.03), which did not persist in the Chr18^A/A^ mice (Supplemental Fig. S5). The velocity of aggregation was significantly reduced in Chr18^A/A^ mice with ADP (p=0.002; Fig. 4A) and significantly reduced in Chr18^+/A^ and Chr18^A/A^ mice with collagen (p=0.0089 and p=0.0002; Fig. 4B). Complete blood counts determined that Chr18^A/A^ mice had significantly lower platelet counts (p=0.02; Fig. 4C) and had increased platelet size variability (p=0.04; Fig. 4D) compared to wildtype littermates, with the Chr18^+/A^ mice having intermediate levels.

**Figure 4:**
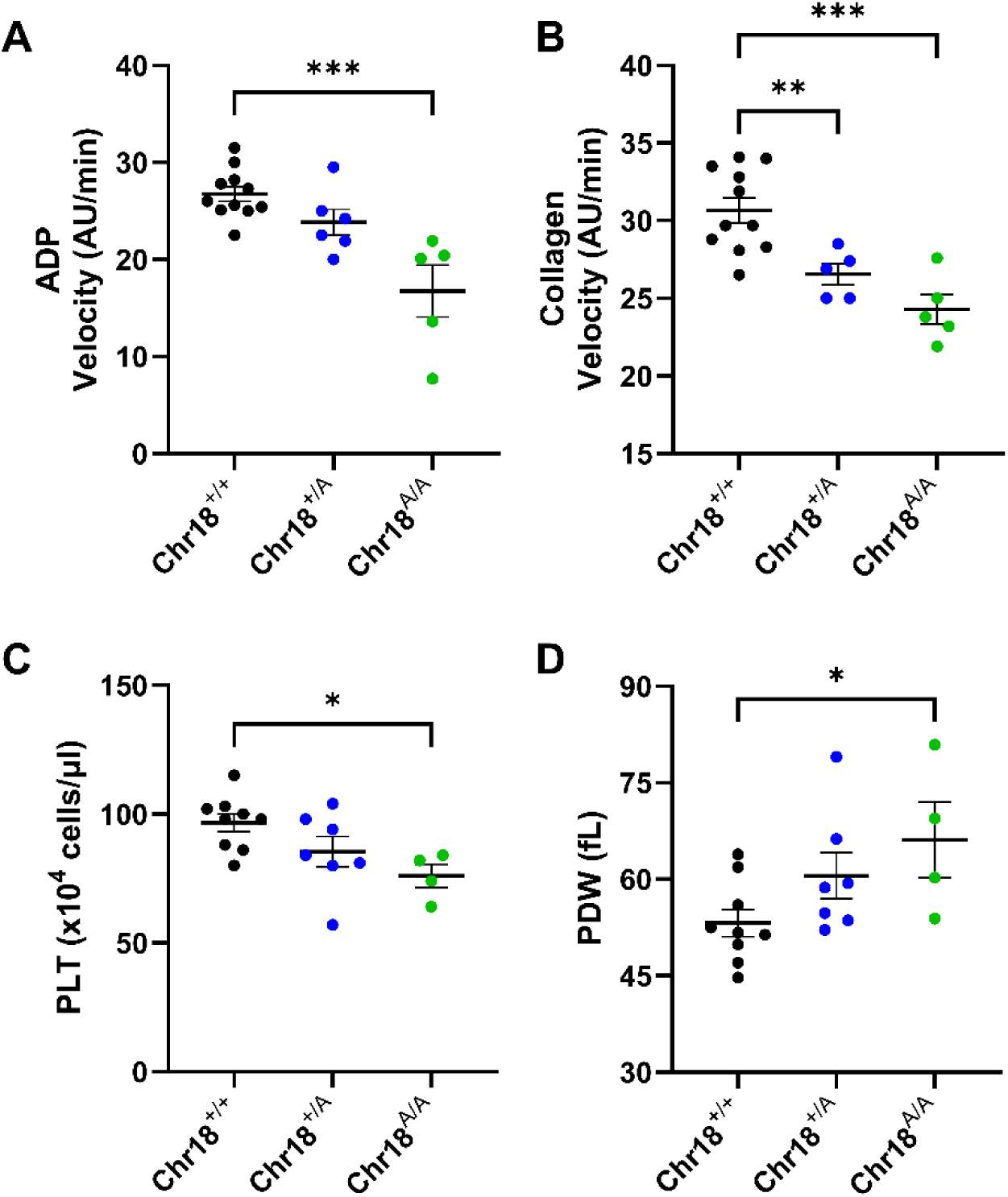
Platelet Aggregation and Blood Counts of Chr18^A^ Mice. **(A)** ADP-induced platelet aggregation velocity. **(B)** Collagen-induced platelet aggregation velocity. **(C)** Platelet (PLT) count. **(D)** Platelet distribution width (PDW). Each data point represents a biological replicate. Mean ± SEM. One-way ANOVA, *p<0.05, **p<0.01, ***p<0.001

In an attempt to identify the cause of this platelet defect, we performed bulk RNA sequencing (RNAseq) on platelets from Chr18^+/+^, Chr18^+/A^, and Chr18^A/A^ mice (n=6). However, based on the principal component analysis (PCA) there was no clear clustering per genotype and no significant differentially expressed genes, suggesting that the altered platelet function may be due to a different mechanism, such as changes at the protein level instead of the transcript level. Taken together, the Chr18^A^ mice have normal coagulation function, but their platelets exhibited a reduced function and slower aggregation kinetics, demonstrating a mechanism by which the Chr18^A^ mutation could be suppressing thrombosis.

### Bulk RNAseq Analysis of Whole Liver from Chr18^A^ Mice Identifies Genes Related to Coagulation and Platelet Activity

To further assess a potential effect of the Chr18^A^ mutation on hepatocytes, as predicted by the DeepSEA algorithm, we performed RNAseq on whole liver samples from Chr18^+/+^, Chr18^+/A^, and Chr18^A/A^ mice (n=6). Although we observed two distinct clusters in the PCA plot, these differences were based on sex rather than genotype (Supplemental Fig. S6A-B). Differential gene expression analysis (based on a p<0.01 and a log_2_ fold change >1 or <-1) between genotypes identified 35 differentially expressed genes (DEGs) between Chr18^+/+^ and Chr18^A/A^ mice, 28 between Chr18^+/+^ and Chr18^+/A^, and 34 DEGs between Chr18^A/+^ and Chr18^A/A^ mice (Supplemental Table S3). Since the biggest effects are expected between Chr18^+/+^ and Chr18^A/A^, we focused on those 35 DEGs. While no differences in classical coagulation genes were found, several genes that have been shown to be directly or indirectly involved in the regulation of coagulation factors or platelet activity were present in this list, which included *Usp2*^29^ (p=0.0051), *Gdf15*^30–33^ (p=0.0024), *Igfbp1*^34^ (p=0.0082), *Pdk4*^35^ (p=0.0023) and *Vim*^36^ (p=0.0034) (Fig. 5).

**Figure 5:**
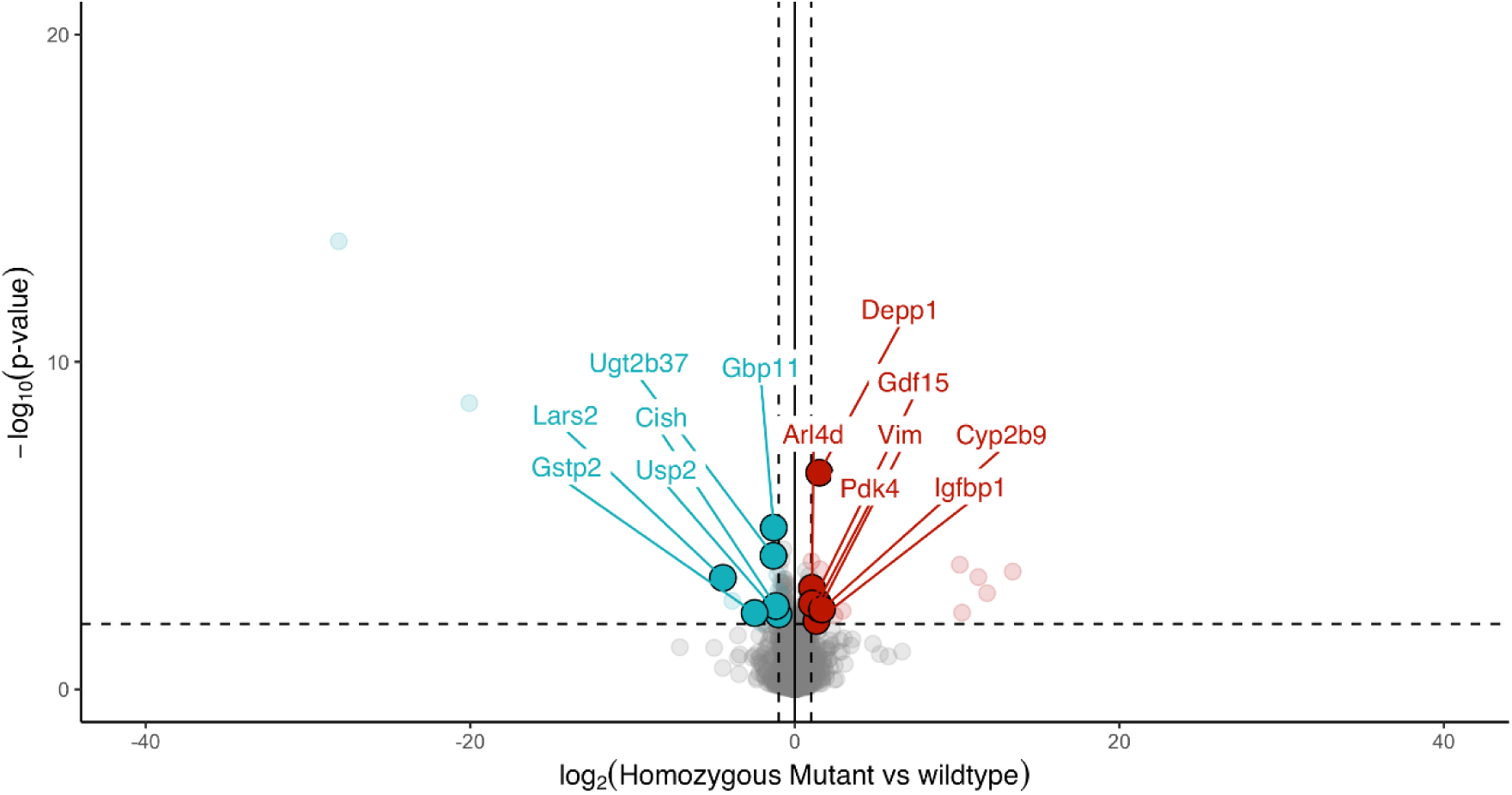
Volcano Plot of Differentially Expressed Gene in Whole Liver Between Wildtype and Chr18^A/A^ Mice. Bulk RNAseq of whole liver from Chr18^+/+^ and Chr18^A/A^ mice (n=6; 3 male and 3 female).

## Discussion

Here we present data from an in-depth whole genome sequencing analysis of one of our large ENU mutagenized mouse *MF5L* suppressor lines (*MF5L16*), where our previous whole exome sequencing analysis was not able to identify a thrombosis suppressor mutation. This line exhibited a biochemical defect in the PT assay, which prompted us to perform whole genome sequencing. This revealed four spontaneously arising suppressor mutations associated with survival of mice carrying the lethal thrombosis *F5*^L/L^ *Tfpi*^+/-^ genotype. The mutation exerting the strongest effect on survival, *sMF5L4*, is an intergenic Chromosome 18 mutation. Importantly, it exerted a significant selective breeding advantage in our *F5*^L/L^ mouse colony and demonstrated a platelet function defect and lead to transcriptional changes in several genes in the liver. The selective breeding advantage of this mutation was further enhanced by inheritance of the other three *sMF5L* mutations. Thus, by superimposing our mutagenesis screen on a B6 genetic background containing antithrombotic mutations selected for during the breeding process, we were able to create the large numbers of *F5*^L/L^ *Tfpi*^+/+^ and *F5*^+/L^ *Tfpi*^+/-^ necessary for conducting our sensitized whole genome ENU mutagenesis screen for identifying dominant thrombosis modifier mutations.

Our sensitized ENU screen for thrombosis modifiers depended on a perinatal lethal phenotype in mice carrying the *F5*^L/L^ *Tfpi*^+/-^ genotype combination. When the *F5*^L/L^ mice were first generated, ∼30% died from spontaneous blood clots and reproduced sporadically^16^. Over the more than 20 generations of backcrosses/intercrosses to produce large numbers of the *F5*^L/L^ genotype necessary to conduct the mutagenesis screen, we noticed fluctuations in the percentage of *F5*^L/L^ mice succumbing to thrombosis. These values ranged from the 30% noted in our initial report, to less than 3%. This is likely explained by the *sMF5L* mutations described here, given our mating strategy of continual backcrossing to the B6 strain followed by intercrosses every∼5 generations to produce *F5*^L/L^ homozygous mice.

We and others have identified spontaneous coding mutations that alter phenotypes^25–28^. Based on our ENU screen, we have previously identified a spontaneous 8 base pair mutation in the *Nbeal2* gene^26^. Homozygous loss-of-function mutations in this gene cause the gray platelet syndrome bleeding phenotype. Interestingly, heterozygosity for this mutation alone does not significantly suppress *F5*^L/L^ *Tfpi*^+/-^ lethal thrombosis. We traced this mutation to a parental 129S1/SvImJ mouse purchased from the Jackson Laboratory, which was used in our ENU mutagenesis cross. This finding prompted us to further investigate the introduced mutations we identified in our mutagenesis screen^27^. We determined that these *sMF5L* mutations selectively spread in our colony until they were robustly represented within the *F5*^L/L^ line, passing to offspring used as breeding stock for our ENU mutagenesis study and bred into several of our ENU lines. Our data suggests that mice carrying these mutations were under positive selection owing to reduced negative effects from *F5*^L/L^ and thus functioned as more robust breeders, producing more offspring and propagating a *F5*^L/L^ line with reduced thrombosis-associated phenotypes.

In conducting whole exome sequencing analysis of a number of our ENU mouse lines that carried a putative thrombosis suppressing mutation, we were surprised to discover that only one of our lines carried a highly penetrant suppressor mutation that altered a structural gene^18^. This observation was inconsistent with the results of other ENU mutagenesis screens, which have demonstrated that the majority of ENU mutants affecting phenotypes were alterations in coding sequences^37^. Nonetheless, this suggested that the thrombosuppressive mutations for many of our lines were within noncoding regions and thus could only be detected by whole genome sequencing. Indeed, using whole genome sequencing to analyze the *MF5L16* ENU line, we successfully identified four novel thrombosuppressive mutations that rescue the lethal phenotype. Unexpectedly, those mutations were not ENU-induced, but spontaneous mutations.

Although the *sMF5L4* mutation resides in a genomic region of unknown significance, we connected this mutation with phenotypic effects in survival, fertility and platelet aggregation, as well as to potential genetic intermediates for those effects via the differential expression of the genes that we discovered through RNAseq of liver samples. While the *sMF5L4* mutation had no effect on the PT, suggesting that one or more of the other *sMF5L* mutations were responsible for influencing the PT, *sMF5L4* rescued *F5*^L/L^ *Tfpi*^+/-^ perinatal lethality at a reduced penetrance (∼15%). This demonstrates that *sMF5L4* alone would not sufficiently account for the entire thrombosuppressive effect in large *MF5L* lines like *MF5L16*. However, as we have shown with the *Actr2* mutation in *MF5L12* and other *MF5L* lines described above, Chr18^A^ and the other three mutations (or also the ENU-induced thrombosuppressive mutations) could work in combination to give rise to numerous *F5*^L/L^ *Tfpi*^+/-^ throughout the lines. Recent evidence has shown that complex diseases, like VTE, are caused by many small effector mutations that work together to contribute to disease^9,10^. The same rationale could hold true for mutations that suppress disease in the presence of disease-causing mutations.

The genomic region surrounding the Chr18^A^ mutation is syntenic to human Chr 18, which contains a lncRNA (ENSG00000299454) within this region. The functionality of this lncRNA is currently unknown but could control genes associated with blood coagulation. Fully characterizing the combinatorial effects of the *sMF5L* mutations in modifying *F5*^L/L^ *Tfpi*^+/-^ survival is of great interest, as these mutations are likely to be true *F5*^L/L^ suppressor mutations judging from how they evolved. Investigating the roles of these mutations in contributing to the suppression of thrombosis should support the data presented here that these mutations - alone or in concert - confer thrombosuppression.

The Chr18^A^ mice have reduced platelet aggregation velocity (AU/min) with both ADP and collagen as agonists, while the area under the curve (AU) and aggregation (U) displayed no significant difference. This suggests the Chr18^A^ platelet defect acts on the rate in which the platelets aggregate but not the maximum aggregation capacity. The Chr18^A^ mice also have reduced platelet counts and increased variability in the platelet distribution width (PDW). Although the comparative bulk transcriptomic analysis of platelet RNAseq was unable to identify a cause for this platelet defect, and additional proteomic analysis might be required to understand the functional defect, we were able to identify suggestive causes for this, such as mutations at the *LTO1* gene locus on chromosome 11 altering platelet counts in humans^38^. In addition, several differentially expressed genes were identified in liver samples, which have been previously associated with platelet reactivity. Of particular interest is the growth differentiation factor 15 (*Gdf15*), which was increased in Chr18^A/A^ mice as compared to Chr18^+/+^ mice. GDF15 has been shown to prevent platelet integrin activation and aggregation, with *Gdf15^-/-^* mice having an accelerated thrombus formation following injury^31^. Moreover, *ex vivo* assays using platelets from deep vein thrombosis patients also indicated that GDF15 could inhibit ADP-induced platelet aggregation^33^, which is in line with our mouse data, and suggests that it could be a potential target for therapeutic interventions in thrombotic disorders. In conjunction with the *sMF5L*, analyzing these mutations and the interactions between them could yield new genomic and mechanistic insights into thrombosis.

Our work highlights an additional layer of complexity arising during the analysis of genetically sensitized ENU screens with a lethal or severe phenotype, since novel spontaneous mutations could arise and modify the sensitizing phenotype itself. In future studies, this could be mitigated by whole genome sequencing each of the breeding stock participants in the pedigree. By discovering that our mutagenesis screen was superimposed on a B6 genetic background containing antithrombotic mutations selected for during the breeding process, we were able to identify additional *MF5L* modifier mutations. These key findings could enable us to pinpoint additional *MF5L* mutants by re-analysis of our previously generated thrombosis suppressor lines. Additionally, these suppressor mutants are critical for understanding the heterogeneity of genetic factors that drive thrombosis in humans.

## Materials and Methods

### Mice

C57BL/6J (B6, stock number 000664) mice were purchased from the Jackson Laboratory. *F5*^L/L^ (*F5*^tm2Dgi^/J, stock number 004080) mice were previously generated^16^. *Tfpi* deficient mice were a generous gift of the late George Broze^39^. The *MF5L* mice were generated in our previous study^19^. The Oakland University Institutional Committee on the Use and Care of Animals and University of Michigan University Committee on the Care and Use of Animals approved all experiments using mice.

### Blood Collection and Plasma Isolation

Male and female mice were anesthetized with sodium pentobarbital and blood was collected by cardiac puncture, with a 1:9 ratio of 3.2% buffered sodium citrate (Aniara) used for an anticoagulant^40^. Whole blood was centrifuged for 10 minutes at 5,000 x g. The plasma was collected and centrifuged again for 10 minutes at 5,000 x g to remove any remaining blood cells. Plasma was stored at-80°C until used.

### Coagulation Assays

Plasma from B6, *F5*^L/L^, *Tfpi*^+/-^, *F5*^+/L^ *Tfpi*^+/-^, *MF5L16 F5*^L/L^ *Tfpi*^+/-^ mice, 8- to 16-week-old Chr18^+/+^, Chr18^+/A^, and Chr18^A/A^ mice (n≥4) was thawed for one minute in a 37°C water bath. The Prothrombin Time (PT) and activated Partial Thromboplastin Time (aPTT) assays were performed on the Delta KC4 (Sigma-Amelung) using RecombiPlastin (Instrumentation Laboratory) and APTT-SP (Instrumentation Laboratory) as previously described^40^.

### FV ELISA

Plasma from 12 Chr18^+/+^, 21 Chr18^+/A^, and 10 Chr18^A/A^ mice was thawed at 37°C and diluted 1:10 and then 1:25 for a final volume of 250μL and a dilution factor of 1:250 in blocking buffer (1X PBS, 0.05% Tween-20, 6% Bovine Serum Albumin solution). Total FV protein concentration was measured by enzyme-linked immunosorbent assay (ELISA). A Nunc Maxisorp plate (Thermo Fisher Scientific) was coated with 50 μL 1 μg/mL GMA-755 (Green Mountain Antibodies) antibody in coating buffer (0.1M Carbonate, pH 9.2) and incubated at 4°C overnight. After washing using 200 μL /well of wash buffer (0.01% Tween-20 in 1X PBS) 3 times, 200 μL of blocking buffer was added and the plate was incubated at room temperature for 2 hours, then washed following the same wash steps as before. Standards and samples were added in duplicate to the plate and incubated at 37°C for 1 hour. A wash was performed and 100 μL of GMA-752 antibody (Green Mountain Antibodies) tagged with HRP (HRP Conjugation Kit-Lightening Link-Abcam) and diluted 1:4000 in blocking buffer for a final concentration of 25 ng/mL, was added to the wells and incubated for 1 hour at 37°C. After washing, 50 μL of TMB was added to the wells in the dark and incubated for 10 minutes at room temperature. The reaction was quenched with 25 μL of 1N H_2_SO_4_ and read immediately on the Biotek Synergy H1 plate reader at 450 nm. Standards were plotted on a log scale, and final measurements were multiplied by the dilution factor^41^.

### Blood Counts and Platelet Aggregation

Citrated whole blood from 8- to 16-week-old Chr18^+/+^, Chr18^+/A^, and Chr18^A/A^ mice was measured on the Advia 2120i Hematology System to determine counts and size and mass parameters of blood cells. The same citrated whole blood was used for type 1 collagen (Hart Biologicals) and ADP (American Biochemical & Pharmaceuticals, LTD)-induced platelet aggregation using the ROCHE Multiplate Aggregometer (Diapharma Group Inc.), following the manufacture’s protocols.

### Tail Bleeding Times

Tail bleeding times were conducted in accordance with the protocol described in Brake *et al*^40^.

### Whole Genome Sequencing

DNA was extracted from tail biopsies using Phenol-Chloroform (Sigma). DNA library preparation and whole genome sequencing were performed by GENEWIZ/Azenta on the Illumina HiSeq X Ten in a 2×150bp paired-end read configuration to obtain ∼30x coverage per sample. The whole genome sequencing data was analyzed by a standard bioinformatics pipeline^42^. The raw reads were aligned to the mouse genome build GRCm38 and genomic positions of candidate variants were updated to and reported as the GRCm39 mouse genome build.

### Sanger Sequencing, Sequence Alignments, and Genotyping

To confirm the WGS candidate mutations, sanger sequencing was performed at the University of Michigan Sequencing Core and as previously described^18^. Briefly, amplicons were generated with the nucleotide of interest using custom outer primer pairs. PCR bands were cut out of a 2% agarose gel and DNA was extracted using the QIAquick Gel Extraction Kit (Qiagen) according to the manufacturer’s protocol. Inner forward and reverse primers were used to bidirectionally Sanger sequence the amplicons. Sequencing chromatograms were visualized using FinchTV (PerkinElmer). The sequences obtained from Sanger sequencing were used to align the DNA sequences from multiple individual mice to each other with MUSCLE, the multiple sequence alignment tool.

DNA for genotyping was isolated from tail biopsies, and mice were genotyped for *F5*^L^ and *Tfpi*^+/−^ as previously described^17^. The introduced mutations on Chr5, Chr9 *4933400C23Rik*, Chr10, Chr13 *Zfp131*, Chr17 *Nrxn1*, and Chr18 *Dcc* were genotyped with a two-reaction multiplex PCR strategy that is sensitive enough to detect wildtype and mutant alleles^18^. Common forward and reverse primers for the were used in separate reactions with a wildtype-specific forward or reverse primer and a mutant-specific forward or reverse primer that differ only in the −1 position on the 3′ end. Inclusion of the common primers provides amplification competition and act as a positive PCR control. The Chr18 intergenic mutation is genotyped by a single set of primers which then undergo a restriction enzyme digest using HinfI (NEB). HinfI cuts the Chr18 mutant sequence (GANTC) while leaving the wildtype sequence (GGNTC) intact. All primers were designed in PrimerBLAST and purchased from Integrated DNA Technologies (IDT) (Supplemental Table S4).

### RNA isolation, sequencing and data analysis

Whole liver samples from 8-15-week-old Chr18^+/+^, Chr18^+/A^, and Chr18^A/A^ mice (n=6; 3 males and 3 females) stored in RNAlater (Sigma) were homogenized in Trizol (Invitrogen) using the gentleMACS tissue dissociator (Miltenyi Biotec). RNA was extracted from the liver samples using the RNeasy Plus Universal Mini Kit (Qiagen) following the manufacturer’s protocol, with an additional Buffer RPE wash step and column DNA elimination step using the RNase-Free DNase Set (Qiagen). RNA was subsequently processed for library preparation using the Kapa Hyperprep kit with RiboErase (Roche), including NEXTflex adapters (Perkin Elmer).

For platelet RNAseq, whole blood was collected via cardiac puncture (described above) from 8-15-week-old Chr18^+/+^, Chr18^+/A^, and Chr18^A/A^ mice (n=6; 3 males and 3 females). Whole blood was mixed with equal volumes of Buffered Saline Glucose Citrate (129 mM NaCl, 13.6 mM Na_3_citrate, 11.1 mM glucose, 1.6 mM KH_2_PO_4_, 8.6 mM NaH_2_PO_4_, pH 7.3) and centrifuged in swinging bucket rotor for 5 mins at 180 x g. The platelet-rich-plasma was isolated and centrifuged in swinging bucket rotor for 10 mins at 1000 x g to pellet the platelets. Trizol was immediately added to the platelet pellet and RNA was extracted using the RNeasy Plus Universal Mini Kit (Qiagen) following the manufacturer’s protocol, with an additional Buffer RPE wash step and column DNA elimination step using the RNase-Free DNase Set (Qiagen). RNA was converted into cDNA and amplified using the SMART-Seq v4 Ultra Low Input RNA kit (Takara Bio USA) according to the manufacturer’s protocol. Full-length cDNA was mechanically sheared using the Covaris E220 Focused Ultrasonicator. Following NEXTflex adapter ligation, libraries were amplified with the Kapa Hyperprep kit (Roche).

Final libraries from both the liver and platelet samples were assessed for quality using the Agilent 4200 TapeStation prior to being sequenced on the Illumina NovaSeq 6000 (MedGenome Inc.) in a 2×100bp paired-end format. Fastq files were quasi-mapped against the mouse reference transcriptome (Mouse Ensembl release 112, GRCm39), followed by transcript quantification via Salmon (PMID: 28263959) (v1.10.0). Transcript counts were loaded into DESeq2 (PMID: 25516281) (v1.44.0) to accommodate differential gene expression analysis, which was performed after excluding low abundant genes (transcript per million mapped reads <1 per group). For the platelet samples, hemoglobin genes were also excluded prior to analysis.

### In silico Regulatory Function

To analyze the functional probability of the introduced mutations, we applied the DeepSEA algorithm for epigenetic prediction of sequence alterations^29^. DeepSEA uses the predictions it makes regarding epigenetic state to calculate z-scores corresponding to the probability that a given sequence is a regulatory feature. A positive z-score indicates that a mutation increases the probability of a regulatory feature and a negative z-score indicates that a mutation decreases the probability of a regulatory feature.

We used 1000 bp sequences with the location of the Chr5, Chr9, Chr13, and Chr18 mutations centered in the middle of the sequence and compared the wildtype sequence and the mutant sequence. The sequences were fed into the DeepSEA framework for functional predictions in hepatocytes, aortic smooth muscle cells, CD34+ mobilized cells, and the megakaryoblastic cancer cell line CMK. The output z-scores from DeepSEA were then converted to percentages and reported here.

### Statistical Data Analysis

Survival analyses were performed with the R *survival* package using *survreg* with Weibull distribution to perform a parametric ANOVA statistical analysis for each variant alone as well as with an interaction term between multiple variants. Survival endpoints were determined when mice perished due to natural causes and mice that were euthanized before that point were censored. The Kaplan-Meier curves were generated with GraphPad Prism^43^. A Student’s t-test was performed to determine significance between the *F5*^L^ breeder pairs. All other data were analyzed in GraphPad Prism with a one-way ANOVA with Dunnett’s multiple comparisons test. Differences were considered significant at p≤0.05, unless otherwise noted.

## Data Access

All whole genome sequencing fastq files generated in this study have been submitted to the NCBI Sequence Read Archive (SRA; https://www.ncbi.nlm.nih.gov/sra/) under accession number PRJNA750698. The RNA-seq reads generated in this study have been submitted to the NCBI Gene Expression Omnibus (GEO; https://www.ncbi.nlm.nih.gov/geo/) (GSEA identifier to follow).

## Acknowledgments

This research was supported by National Institutes of Health, National Heart, Lung, and Blood Institute (NHLBI) Grants R15-HL133907, R15HL172202 and R01-HL135035, and an American Heart Association (AHA) Innovative Research Grant 17IRG33460238 and AIREA Research Grant 23AIREA1055261 (to R.J.W.). M.A.B was supported by the AHA MWA Undergraduate Student Research Grant 15UFEL25700204. C.D.S. and K.H.M were supported by the AHA Summer Undergraduate Research Program, 23IAUST1034173. A.M.J. was supported by the M-KUHR training grant, TL1DK136046, and the NSF NRT grant 2152007. We thank David Ginsburg for his long-term support of this project and critical reading of this manuscript.

## Authorship Contributions

M.A.B., A.C.C., S.T., M.A.vE., G.Z., J.K., T.M.P., C.D.S., A.M.J., K.H.M., A.E.S., and R.J.W. performed experiments; M.A.B., A.E.S., A.C.C., M.A.vE., J.K., T.M.P., C.D.S., A.M.J., K.H.M., F.U.B., A.C.C, D.R.S., A.E.T., D.R.B., and R.J.W. analyzed results and made the figures; M.A.B., A.C.C., M.A.vE., J.K., C.D.S., A.M.J., A.E.T., D.R.B., and R.J.W. designed the research and wrote the paper.

## Disclosure of Conflicts of Interest

The authors declare no competing financial interests.

## Notes

### Competing Interest Statement

The authors have declared no competing interest.

