## Supplemental Figures for "A Sensitized ENU Mutagenesis Screen for Thrombosis Modifiers Identifies Suppressor Variants in Non-mutagenized Parental Generations Due to Antithrombotic Selective Pressures"

**A**

**Chr12 IR1**
**C→T**
**12:105436177**

```
NCBI_Ref_B6       CCCATCCCACCAGCCCAATGCAGTGTACAC------------------------------
Sequenced_B6      CCCATCCCACCAGCCCAATGCAGTGTACACAGAGAGTCCTCCACCCATCCCACCAGCCCA
                  ******************************

NCBI_Ref_B6       ------------------------------------------------------AGAGAG
Sequenced_B6      ATGCAGTACACAGAGAGTCCTCCACCCATCCCACCAGCCCAATGCAGTGTACATAGAGAG
                                                                        ******
```

**B**

**Chr16 IR2**
**T→G**
**16:40539588**

```
Sequenced_B6      AATTTCCNAGAGAGAGAGAGAGAGNGNGAGAGA--------------GAGAGAGAG----
Sequenced_129S1   AATTGCAGAAAGAGAGAGAGAGAGAGAGAATGAATCCTTGTCCACTTGATGCAGAAGTCC
                  **** *  * ************** * **  **              **   ***

Sequenced_B6      ------AGAGGGGGGGGGGGTAAGGACCCTCAGCATCTCAGCTCTATGCTGTGATGGTTT
Sequenced_129S1   ATGTGTATACTGTGTCCAGGTAAGGGCCCTCAGCATCTCAGCTCTATGCTGTGATGATTT
                        * *  * *    ******* ****************************** ***
```

**C**

**Chr18 IR3**
**T→C**
**18:82164154**

```
Sequenced_B6      CTAGGGCAGGGCTCCCCTCTGTCTCCTCATCACCAGCTCACAGCGCCAGGGCAACCCCAC
Sequenced_129S1   CTAGGGGCGGGCTCCCCTCTGTCTCCTCATCACCAGCTCACAGC----------------
                  ******  ************************************

Sequenced_B6      TATCTCCTCATCACCAGCTCACAGCACCAGGGCAACCCCACTATCTCATCACCAGCTCAC
Sequenced_129S1   ------------------------------------------------------------

Sequenced_B6      AGTGCCAGGGCAACACCTCTGTCTCCTCATCACCAGCTTACAGTGCCAGAGCAACCCCTC
Sequenced_129S1   --------------------------------------------GCCAGAGCAACCCCTC
                                                              ****************
```

**Figure S1: Uncharacterized B6 and 129S1 Genomic Indels.** The red letters represent the nucleotide called as a mutation. **(A)** Intergenic region on chromosome 12. The grey highlighted regions are the repeated sequence. The blue letter represents the nucleotide believed to be calling the heterozygous mutation. **(B)** Intergenic region on chromosome 16. The purple lines represent start and end of sequence variation. The grey highlighted region is the mononucleotide repeat of Gs. **(C)** Intergenic region on chromosome 18. The blue letters represent the nucleotide believed to be calling the heterozygous mutation. The grey highlighted regions are the repeated sequence.

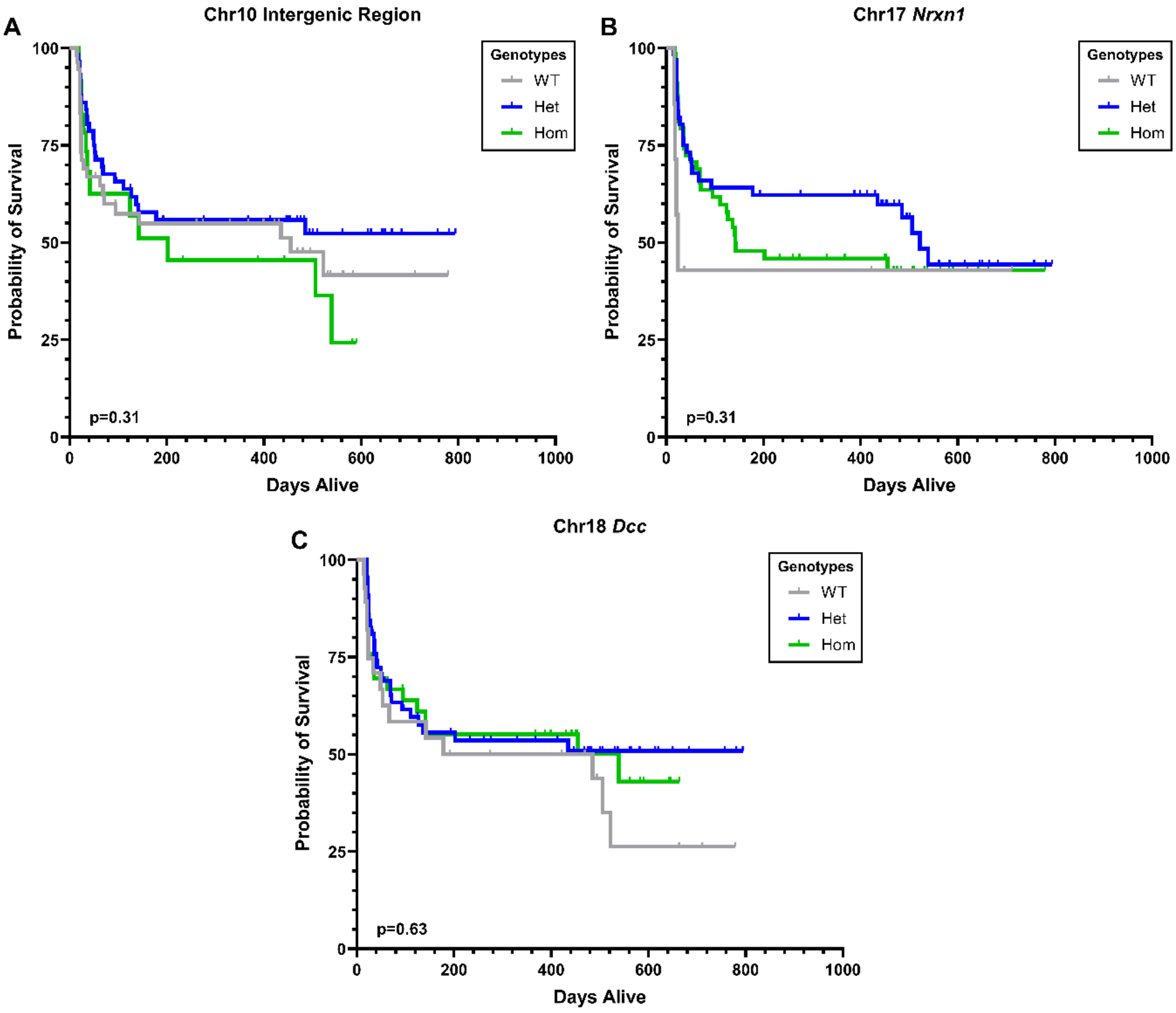


**Figure S2: Nonsignificant Introduced Mutations.** Kaplan-Meier survival curves of *F5*$^{L/L}$ *Tfpi*$^{+/-}$ mice from *MF5L16*. **(A)** Curve between WT (n=55), Het (n=57), and Hom (n=24) for the Chr10 intergenic region mutation. **(B)** Curve between WT (n=7), Het (n=64), and Hom (n=66) for the Chr17 intronic mutation in *Nrxn1*. **(C)** Curve between WT (n=28), Het (n=67), and Hom (n=42) for the Chr18 intronic mutation in *Dcc*. WT=wildtype; Het=heterozygous for mutation; Mut=homozygous for mutation

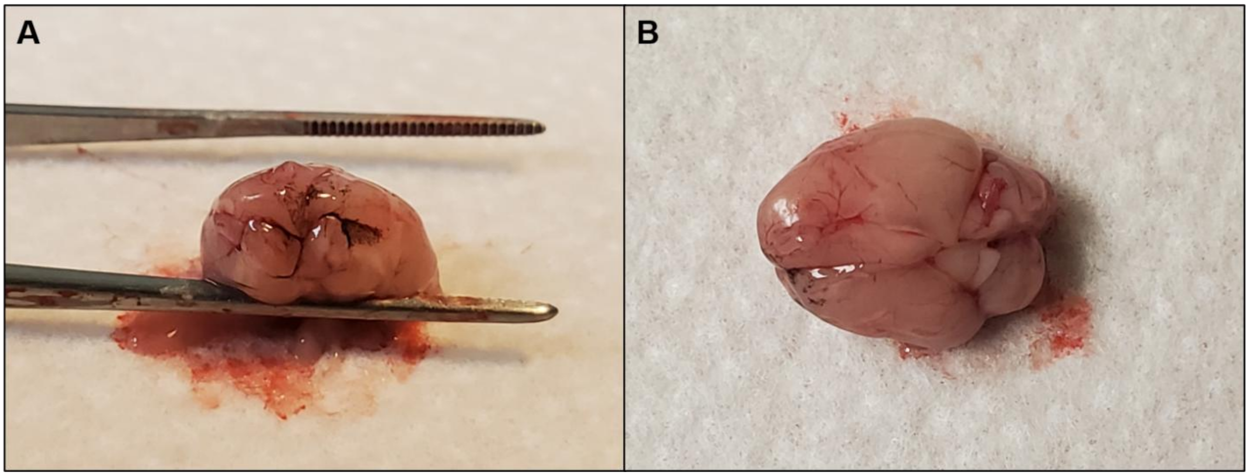


**Figure S3: Brain of *F5*$^{L/L}$ *Tfpi*$^{+/-}$ Chr18$^{+/A}$ Male Mouse with Constant Seizures. (A)** Blood clots in the frontal lobes of the brain. **(B)** A shrunken left lobe of the brain.

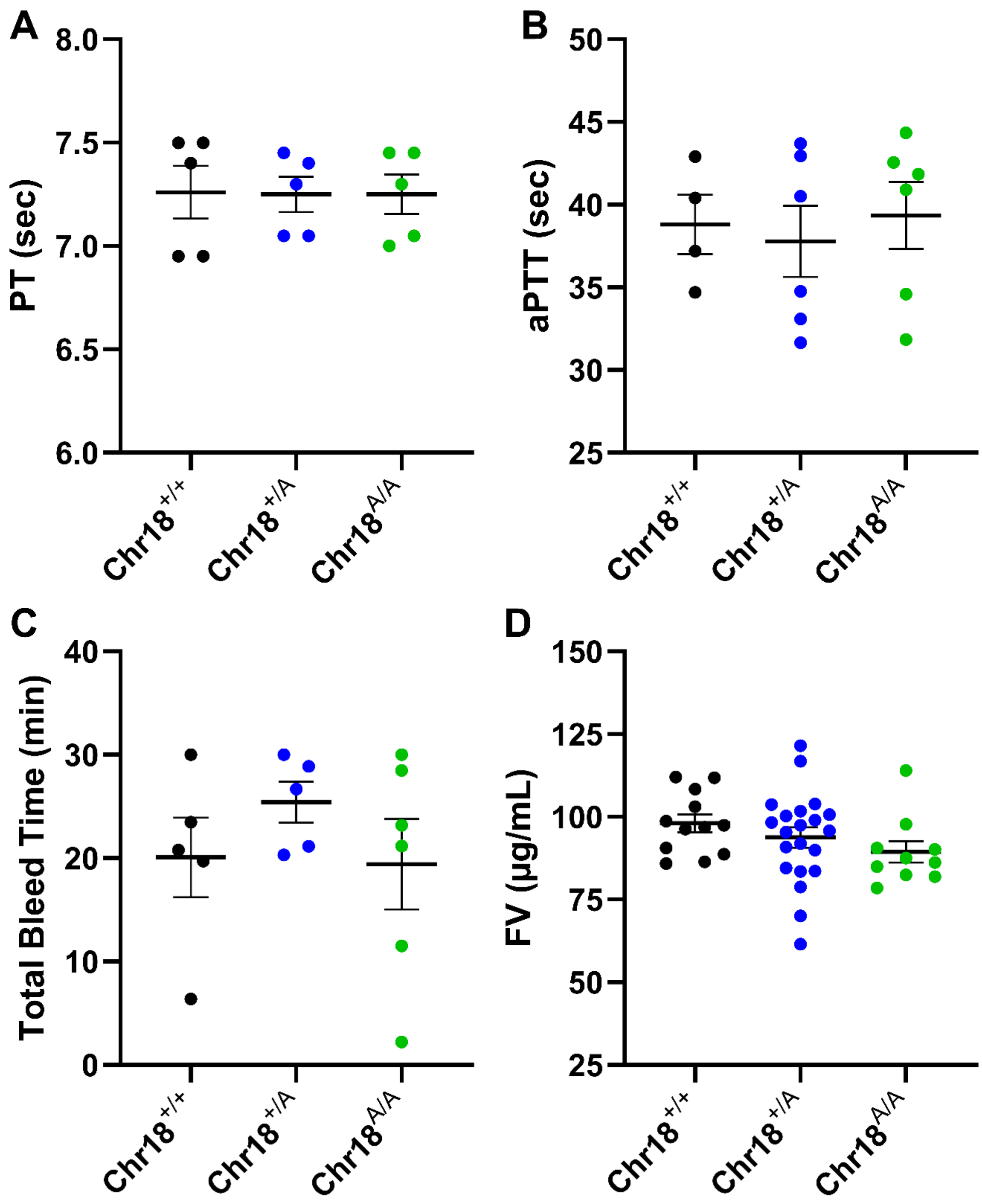


**Figure S4: Coagulation Assays, Tail Bleed, and FV levels of Chr18$^{A}$ Mice. (A)** The Prothrombin Time (PT) and **(B)** activated Partial Thromboplatin Time (aPTT) coagulation assays. **(C)** Total bleed time after tail transection. **(D)** Plasma FV levels measured by ELISA. Each data point represents a biological replicate. Mean ± SEM.

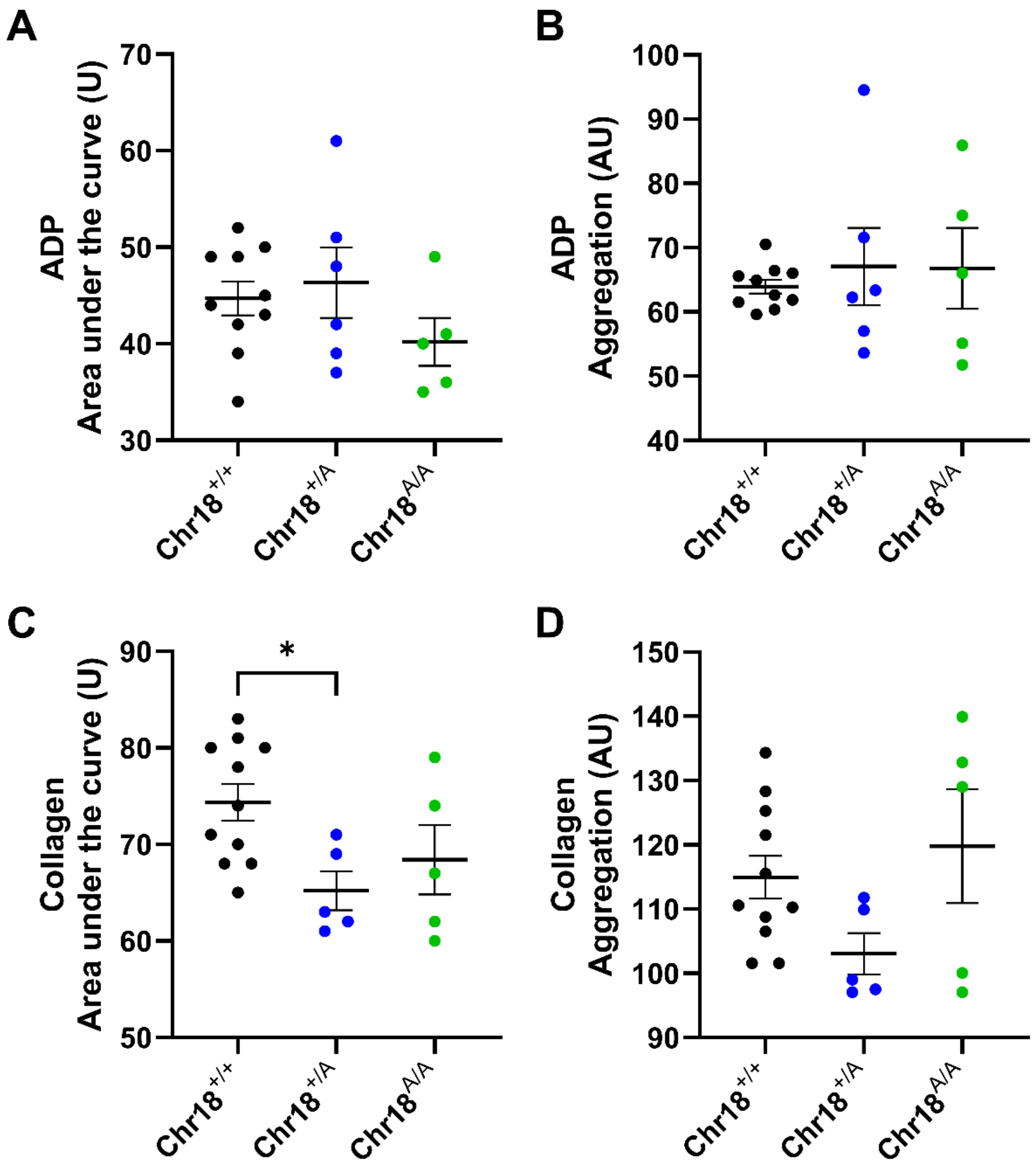


**Figure S5: Platelet Aggregation. (A)** Area under the curve (U) and **(B)** Aggregation (AU) of ADP-induced platelet aggregation. **(C)** Area under the curve (U) and **(D)** Aggregation (AU) of collagen-induced platelet aggregation. Each data point represents a biological replicate. Mean ± SEM. One-way ANOVA, *p<0.05

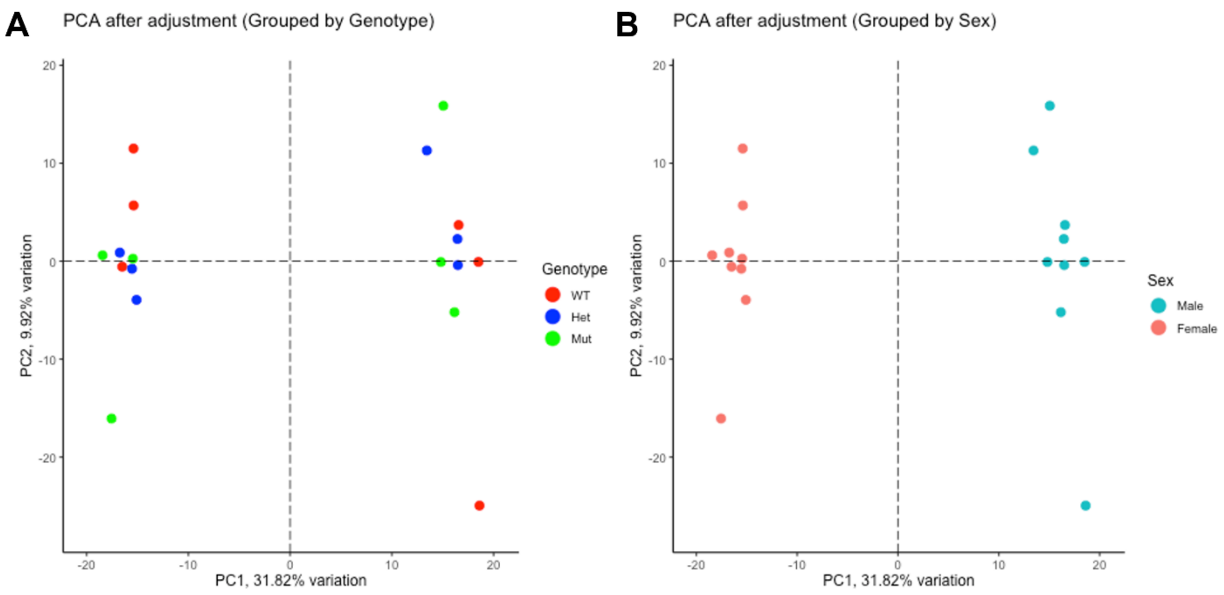


**Figure S6: Principal Component Analysis (PCA) Plots of Bulk Liver RNAseq. (A)** PCA plot from whole liver RNAseq analysis to visualize the transcriptional relationship between Chr18$^{+/+}$ (WT), Chr18$^{+/A}$ (Het), and Chr18$^{A/A}$ (Mut) mice (n=6). **(B)** PCA plot from whole liver RNAseq analysis to visualize the transcriptional relationship between male and female mice (n=6). Each data point represents a biological replicate.
